# Genome-wide identification of protein binding sites in mammalian cells

**DOI:** 10.1101/2021.09.29.462333

**Authors:** Fenglin Liu, Tianyu Ma, Yu-Xiang Zhang

**Author notes:** Yu-Xiang Zhang (corresponding author). **Email addresses** Fenglin Liu (first author) Tianyu Ma. **Funding** This work was funded by grants from Beijing Municipal Natural Science Foundation (grant number 7192022) and the National Natural Science Foundation of China (NSFC; grant number 81372156).

## Abstract

We present GWPBS-Cap, a method to capture genome-wide protein binding sites (PBSs) without using antibodies. Using this technique, we identified many protein binding sites with different binding strengths between proteins and DNA. The PBSs can be useful to predict transcription binding sites and the co-localization of multiple transcription factors in the genome. The results also revealed that active promoters contained more protein binding sites with lower NaCl tolerances. Taken together, GWPBS-Cap can be used to efficiently identify protein binding sites and reveal genome-wide landscape of DNA-protein interactions.

## Introduction

DNA-binding proteins regulate gene expressions through binding to specific DNA sequences. With the development of high throughput sequencing technology, more and more genome-wide protein binding sites (PBSs) have been obtained by a series of biological methods. For example, the chromatin immunoprecipitation (ChIP) has been widely used in this field [1-2]. However, there are shortcomings in this method. Firstly, specific antibody for each target protein is required, and some proteins cannot be tested due to lack of suitable antibodies; Secondly, a one-time ChIP experiment can only provide the binding profile for one protein. Indeed, effective capturing the genome-wide precise PBSs is needed.

Here we describe a method, called genome-wide protein binding site capture (GWPBS-Cap), which can be used to map genome-wide protein binding sites (PBSs) without using antibodies. In the procedure, DNA and DNA-binding proteins are cross-linked with formaldehyde, and the cross-linked chromatin is sonicated and conjugated to magnetic beads using Sulfo-SMCC. Combining with high throughput sequencing, Genome-wide protein binding sites can be identified.

The DNA-protein interactions *in vivo* are closely related to binding strength. Given that protein-DNA interactions are largely electrostatic in nature and are sensitive to buffers with high ionic strength (high salt concentration), increased salt concentration adversely affects protein binding to DNA, and DNA-protein interactions vary in their sensitivities to salt concentration [3-4]. Washing nuclei with buffers containing different concentration of NaCl is known to result in the isolation of dramatically different chromatin fractions [5]. In the GWPBS-Cap assay, the nuclei are treated with washing buffers containing various concentrations of NaCl, the interacting DNAs and proteins are cross-linked, and the protein binding sites are then identified. The dynamics of DNA-proteins interactions can then be obtained. Our results show that GWPBS-Cap assay can be used to capture the genome-wide PBSs and will be a useful tool for understanding the landscape of DNA-protein interactions. GWPBS-Cap will be an effective complementary tool to ChIP-Seq.

## Results

### The GWPBS-Cap flowchart and definition of PBSs

We have developed GWPBS-Cap, a method designed for genome-wide capture of protein binding sites without using antibodies (Figure 1A, Materials and methods). To capture cross-linked chromatin complexes, we used Sulfo-SMCC (sulfosuccinimidyl 4-[N-maleimidomethyl]cyclohexane-1-carboxylate) to conjugate amine groups on the magnetic beads and the sulfydryls on the chromatin proteins (Figure S1). The amine groups on the magnetic beads were first activated with Sulfo-SMCC. Prior to the conjugation with the pre-activated beads, sheared chromatin was incubated with TCEP (tris(2-carboxyethyl)phosphine) in order to reduce the disulfide bonds of proteins to sulfydryls. Once bead-bound, the DNA fragments in the captured chromatin complexes were subjected to additional chemical processing steps, such as 5’-3’ cutting by lambda exonuclease [6] to improve the resolution of the binding sites. Adapters were then added to the DNA fragments and used for amplification by PCR. The fragments in the resulting DNA libraries ranged from 80-120 bp and were subjected to high-throughput sequencing. Reverse complementary regions of pairs of overlapping 5’ end-sequencing reads were defined as protein binding sites (PBSs).

**Figure 1.**
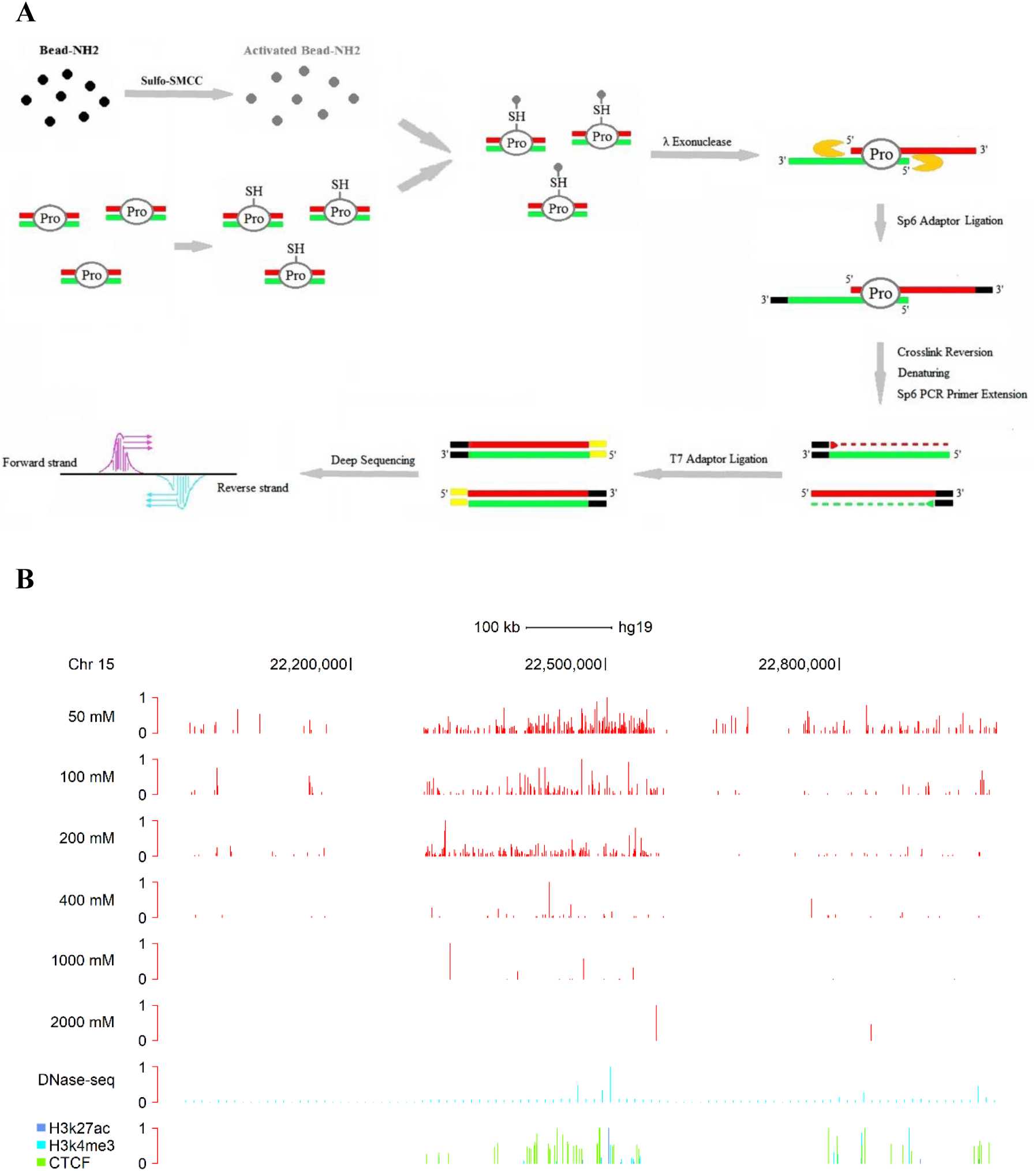
Identification of genome-wide protein binding sites with different binding strengths. **(A)** A schematic diagram of GWPBS-Cap in this study. The proteins from cross-linked chromatin complexes are covalently conjugated with amine-containing beads through Sulfo-SMCC. The bound DNA fragments are digested with 5’ to 3’ exonuclease (lambda exonuclease), which trims the 5’ ends of DNA up to the site of cross-linking. After high-throughput sequencing, the 5’ ends of the sequencing reads are mapped to the reference genome. Reverse-complementary region of a pair of 5’ end reads is considered a protein-DNA cross-linking site and a PBS. (**B)** Comparison of GWPBS-Cap to ChIP-seq and DNase-seq assays at a locus on chromosome 15 in Hela cells. PBSs were obtained using nuclei washed with buffers containing six different concentration (50 mM, 100 mM, 200 mM, 400 mM, 1000 mM, 2000 mM) of NaCl and their PBS peaks are shown. DNase-seq (Duke) data was from obtained from the ENCODE Consortium datasets. Bottom, composite locations of CTCF and histone modifications associated with active enhancers and promoters. PBS, protein binding site; Pro, protein.

Different types of DNA-binding proteins may bind to DNA with different strengths. To test DNA-proteins binding strengths, we incubated nuclei in buffers with different ionic strengths (different salt concentration) before the DNA-protein to be cross-linked. In order to remove proteins with different DNA-binding strengths, nuclei were washed with buffers containing six different concentrations (50 mM, 100 mM, 200 mM, 400 mM, 1000 mM, 2000 mM) of NaCl, respectively, yielding fractions 1 to 6. After removing sensitive DNA binding proteins using above salt buffers, the chromatin fractions collected were cross-linked with formaldehyde, yielding chromatin samples 1 to 6, which were then sonicated and conjugated to the magnetic beads with Sulfo-SMCC. DNA adapters were ligated to the DNA fragments on the beads. The DNA fragments were then extracted, and DNA libraries were prepared for each sample and used in next generation sequencing. Six sets of high throughput sequencing data were thus obtained.

On average, about 186,700,000 unique GWPBS-Cap sequence reads were obtained from each set. We found that some PBSs show up in different groups. For example, the PBS in 400 mM salt group which located on 121481274-121481356 bp, Chromosome 1, overlaps with three other PBSs from 50 mM, 100 mM and 200 mM groups (Figure S2A). As these four PBSs overlap each other, we regarded these four PBSs as the same PBS. We then classified these PBSs according to the maximum salt concentration that they tolerate. For example, if a given PBS was shown up in the 50 mM and 100 mM groups, then 100 mM NaCl was the maximum salt concentration it could tolerate, and the PBS is considered to belong to the 100 mM NaCl group (Figure S2B). According to the above classification criteria, 842,855, 367,415, 548,381, 114,691, 20,432 and 698 PBSs were obtained in 50 mM, 100 mM, 200 mM, 400 mM, 1000 mM and 2000 mM NaCl groups, respectively (NCBI, GEO, GSE116770_PBS-peaks_of_the_six_NaCl_groups.xlsx). We compared data from six groups to DNase-seq data, and found that GWPBS-Cap-seq had a signal-to-noise ratio similar to that of DNase-seq from the ENCODE (Encyclopedia of DNA Elements) Consortium (Figure 1B). At a locus previously highlighted by others, the density of PBSs decreased as the concentration of NaCl in the washing buffer increased (Figure 1B and Figure S2C).

### Identification of nucleic proteins eluted with solutions containing different concentrations of salts

Sodium ion concentration affects DNA-protein binding, so if a DNA-protein binding can withstand high sodium ion concentration, it indicates that the DNA-protein binding force is stronger. To examine the binding affinity between different nucleic proteins and DNA, we collected several protein fractions eluted with solutions containing different concentrations (50 mM, 50-100 mM, 100-200 mM, 200-400 mM, 400-1000 mM) of NaCl. For example, “50 mM” means that the nuclear proteins were extracted with 50 mM salt. “50-100 mM” means that the nuclei were washed with 50 mM salt first and then 100 mM salt was used to extract the nuclear proteins. Mass spectra were used to identify proteins in the different fractions. The results are shown in Table S1. Most of the groups were enriched for proteins with Gene Ontology (GO) molecular function terms, but the frequencies of the different GO types in the five fractions were different (Figure S3). We analyzed the distribution of transcription factors (TFs), chromatin remodeling proteins, histones, histone variants and histone modifiers in the different elution groups (Figure 2A). The contents of histone and histone variants identified by mass spectrometry increased significantly with the increase of NaCl concentration. The elution of the other three types of proteins showed a parabolic distribution, with the highest content in the 100-200mM NaCl fraction. The results showed that 100-200mM NaCl was the optimal elution condition for TFs and other regulatory proteins. The histone modifiers were sensitive to NaCl concentration and almost zero in the elution group of 1000mM NaCl. To our surprise, there were still many TF proteins eluted in the 1000mM NaCl elution group, indicating some TFs or TF complexes have very high binding strength to DNA.

**Figure 2.**
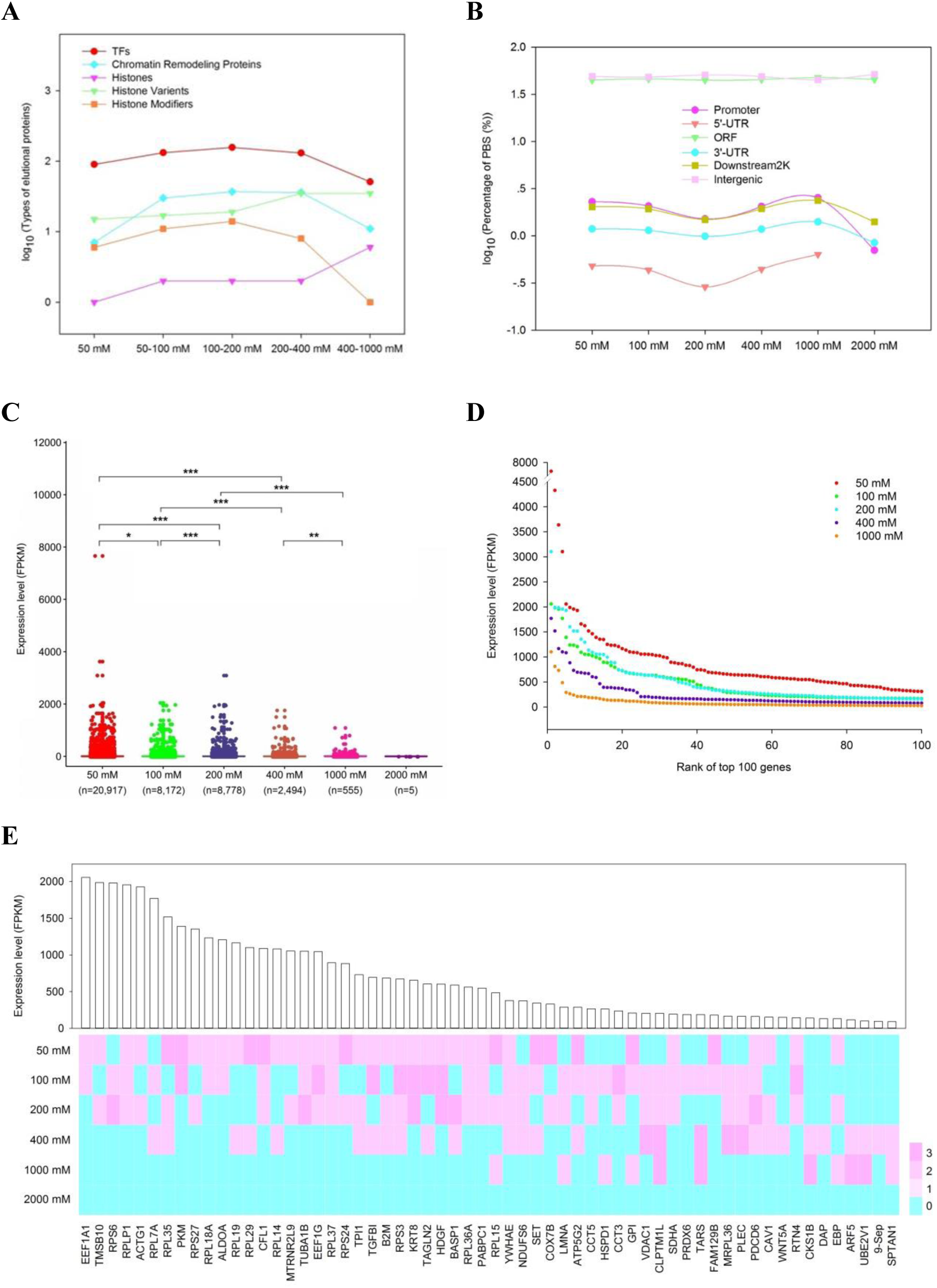
Distribution of protein binding sites (PBSs) in functional gene regions. **(A)** Distribution of proteins in five GO categories (TFs, Chromatin remodeling proteins, Histones, Histone variants and Histone modifiers) in the successive salt buffer elution groups. **(B)** Percentages of PBSs associated with different gene regions, including promoters, UTRs, ORFs, downstream regions and intergenic regions. **(C)** Overall correlation between gene expression level and PBS distribution pattern in promoters. The distribution pattern of PBSs in 6 different salt groups are shown. The gene expression level for each gene was computed with normalized FPKM. The data were analyzed for statistical significance between two different groups using the Cruskal-Wallis test. *P < 0.05, **P < 0.01, ***P < 0.001. **(D)** Top 100 genes were selected for each group of five PBS groups, according to the number of PBSs contained in gene promoters. Horizontal axis represents the ranks of top 100 genes in each PBS salt group. Vertical axis shows the expression level of the genes. **(E)** The distribution pattern of PBSs in the promoters of 60 random selected genes. Top panel shows the FPKM level of each gene. Bottom panel uses different colors to indicate numbers of PBSs in each PBS group. PBSs in the promoters of highly expressed genes tend to be enriched in the lower salt groups.

### The distribution of PBSs in different gene regions

We also analyzed the distribution for PBSs from the six NaCl groups in different gene regions (Figure 2B). The distribution of PBSs in promoters, UTRs, and downstream (2 kb) regions followed parabolic curves. When the concentration of NaCl in the washing solution was less than 200 mM, the number of PBSs in the in these gene elements or downstream regions was negatively correlated with the concentration. When the concentration of NaCl in the washing solution was more than 200 mM, the number of PBSs in these gene elements or downstream regions was positively correlated with the concentration of NaCl. This phenomenon was especially reflected in promoter PBSs, suggesting that protein-DNA interaction in the promoter region is very dynamic. In ORF and intergenic regions, the distributions of PBSs in different Na groups are not significantly different, suggesting that these regions bind to a variety of different types of proteins.

### Gene expression levels and PBS distribution in the gene promoters

To further investigate the relationship between PBS regions with different binding strengths and expression levels of corresponding genes, RNA-seq experiments were performed. The fragments per kilobase of exon per million fragments mapped (FPKM) of genes in Hela cells was calculated (Table S2).

We evaluated the correlation between gene expression level and the distribution of PBSs in 25,607 promoters. FPKM data were calculated and their correlations with PBSs in their promoters were explored (Table S3). We found that the binding strength of PBSs to promoters was inversely correlated with gene expression level (Figure 2C). We selected top 100 genes for each of five PBS groups, according to the number of each group PBSs contained in gene promoters. The genes whose promoters contain more PBSs tend to be highly expressed (Figure 2D). To gain further insight into the correlation between gene expression and PBSs, we compared the distributions of PBSs in the promoters of the 50 most highly expressed genes and 50 non-expressed genes (Figure S4). In the highly expressed genes, most PBSs in their promoters were in the lower salt groups, and none were in the 1000 mM or 2000 mM salt groups, indicating that DNA binding proteins with weak binding strengths are more likely to bind to the promoters of highly expressed genes. Nevertheless, silenced genes also had some PBSs in the lower salt groups in their promoters, indicating that the promoters of silenced genes also have PBSs and display dynamic protein-DNA interactions. We next randomly selected 60 genes and calculated the number of PBSs in their promoters (Figure 2E), and we confirmed that the PBSs in the promoters of genes with higher FPKM values were enriched in the lower salt groups.

### Prediction of TFBSs from PBSs and their base compositions

In order to predict which proteins bind to PBSs and regulate gene expression, we focused on TFs, a large class of DNA sequence-specific binding proteins with regulatory functions. We mapped our PBSs to the human transcription factor binding site (TFBS) datasets from the ENCODE Consortium [7-8]. Every TFBS that falls into PBS was considered to have binding site in a PBS. About 96.67% of TFBSs provided by the ENCODE Consortium, could be found in the PBSs, indicating that GWPBS-Cap is an effective method for capturing protein binding sites (NCBI, GEO, GSE116770_Predictions_for_the_six_groups_of_TFBSs_located_in_PBSs.xlsx). The TFs that predicted to bind to these PBSs fall into 26 classes [9-10]. The genome-wide representation of the TF binding profile is shown in Figure 3A-D. The results are shown in order of the number of binding sites of these TF families, suggest that TF families with many more genome-wide binding sites have a similar genome-wide distribution profile.

**Figure 3.**
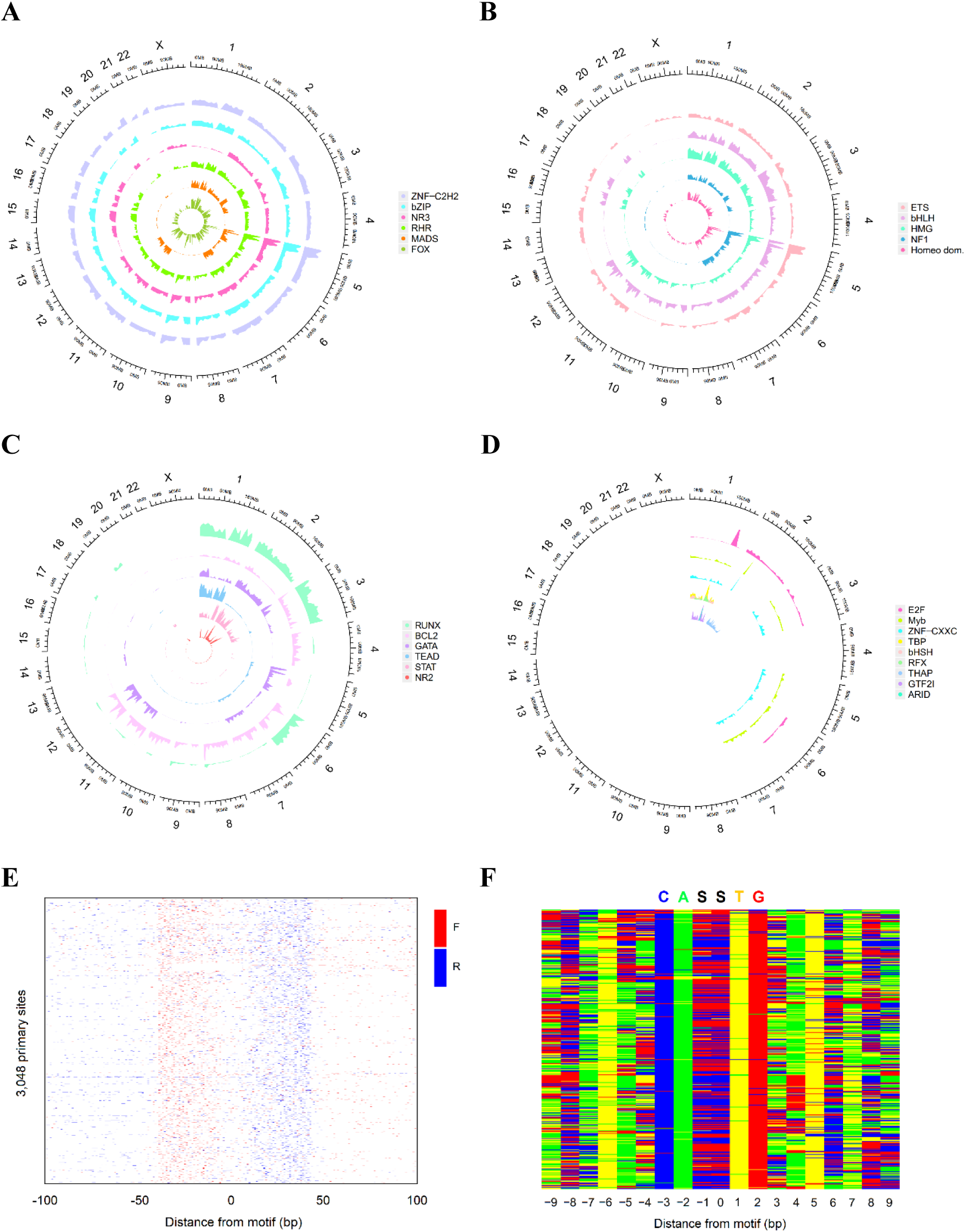
TFBSs were predicated from PBSs identified using GWPBS-Cap. **(A*-*D)** Circos plots of distribution of binding sits of different transcription factor (TF) families on 23 chromosomes. The results are shown according to the number of genome-wide binding sites of different TF families. From figure A to figure D, the binding sites of different TF families in the genome are shown in descending order. From outer track to inner track, the number of genome-wide binding sites of different TF families are shown in descending order in each circos-plot. **(E)** Raw sequencing tag distribution of predicated 3,048 primary TCF3-bound locations (rows). Red and blue represent the 5’ ends of forward (left border) and reverse strand tags (right border), respectively, centered by the motif midpoint. **(F)** Color chart representation of the TCF3 consensus sequence located in the middle of peak pair of the 5’ ends of forward and reverse strand tags. Each row represents a bound sequence ordered as in figure E. Green, yellow, red and blue indicates A, T, G, and C. The TCF3 consensus sequence is indicated as GASSTG (S=C/G).

We analyzed the base compositions of some TFs from the TFBS collection. For example, TCF3 factors have a well-defined DNA recognition site (CASSTG) that could be used for independent validation. We randomly selected 3,048 PBSs which are predicated to bind with TCF3 (Table S4) and found that their related PBS tags located on two strands have exonuclease barriers (Figure 3E). The sequences between the barrier pairs contained the TCF3 recognition site CASSTG (Figure 3F). This indicates that indeed TFBSs are captured by GWPBS-Cap.

To confirm that our predicted TFBSs are real TF binding sequences, we used SP1-ChIP to test our predictions. We selected one SP1 TFBS each in the 100 mM and 1000 mM salt groups (Materials and methods), and carried out ChIP-PCR to confirm the existence of these two sites (Figure S5). The result means that the PBSs captured by GWPBS-Cap can be used to predict TFBSs.

### Sequence features of PBSs containing multiple TFBSs

As it has been reported that it is common for multiple TF families to bind to the same short region of the genome [11-12], we found that almost PBSs are predicted to contain binding motifs of multiple TF families. This suggests that different TF-families may interact in the same DNA region. We listed the top 20 TF-family pairs which motifs appear on the same PBS in Table 1.

**Table 1.** The top 20 transcription factor-family pairs binding on the same PBSs.

| **Rank** | **TF-family pairs** | **Number** |
| --- | --- | --- |
| **1** | ZNF-C2H2 – bZIP | 1,579,447 |
| **2** | ZNF-C2H2 – NR3 | 1,248,235 |
| **3** | ZNF-C2H2 – RHR | 1,242,105 |
| **4** | bZIP – RHR | 1,170,617 |
| **5** | bZIP – NR3 | 1,159,501 |
| **6** | ZNF-C2H2 – FOX | 1,028,604 |
| **7** | bZIP – FOX | 978,942 |
| **8** | ZNF-C2H2 – ETS | 964,872 |
| **9** | NR3 – RHR | 945,050 |
| **10** | bZIP – ETS | 922,509 |
| **11** | FOX – RHR | 806,498 |
| **12** | ETS – NR3 | 806,383 |
| **13** | ETS – RHR | 788,157 |
| **14** | ZNF-C2H2 – bHLH | 783,526 |
| **15** | FOX – NR3 | 759,936 |
| **16** | bHLH – bZIP | 736,260 |
| **17** | ETS – FOX | 688,364 |
| **18** | bHLH – RHR | 448,656 |
| **19** | bHLH – NR3 | 446,308 |
| **20** | bHLH – FOX | 293,099 |

To explore the potential protein-protein interactions between different TF families that bind to the same PBS. We selected two TF-pairs, NFIC-TCF12 and RUNX3-EGR1, which are derived from different TF-families, respectively. We randomly selected 272 PBSs containing both NFIC and TCF12 binding sites (Table S4) and 100 PBSs containing both RUNX3 and EGR1 binding sites (Table S4). These PBSs with multiple TFBSs also present obvious peak pairs of exonuclease barriers (Figure 4A-B). We used MEME Suite [13] to find motifs from these PBSs and compared them with their single motifs downloaded from JASPAR website [14]. We found that the motifs in the PBS combining two TFBSs is the overlapping two motifs corresponding to the two TF recognition sites (Figure 4C-D), e.g. the sequences of NFIC-TCF12 recognition sites much like the two partially overlapped TF recognition sequences (Figure 4C). This is consistent as previously reported [11]. Thus, our findings are helpful for predicting protein-protein interactions.

**Figure 4.**
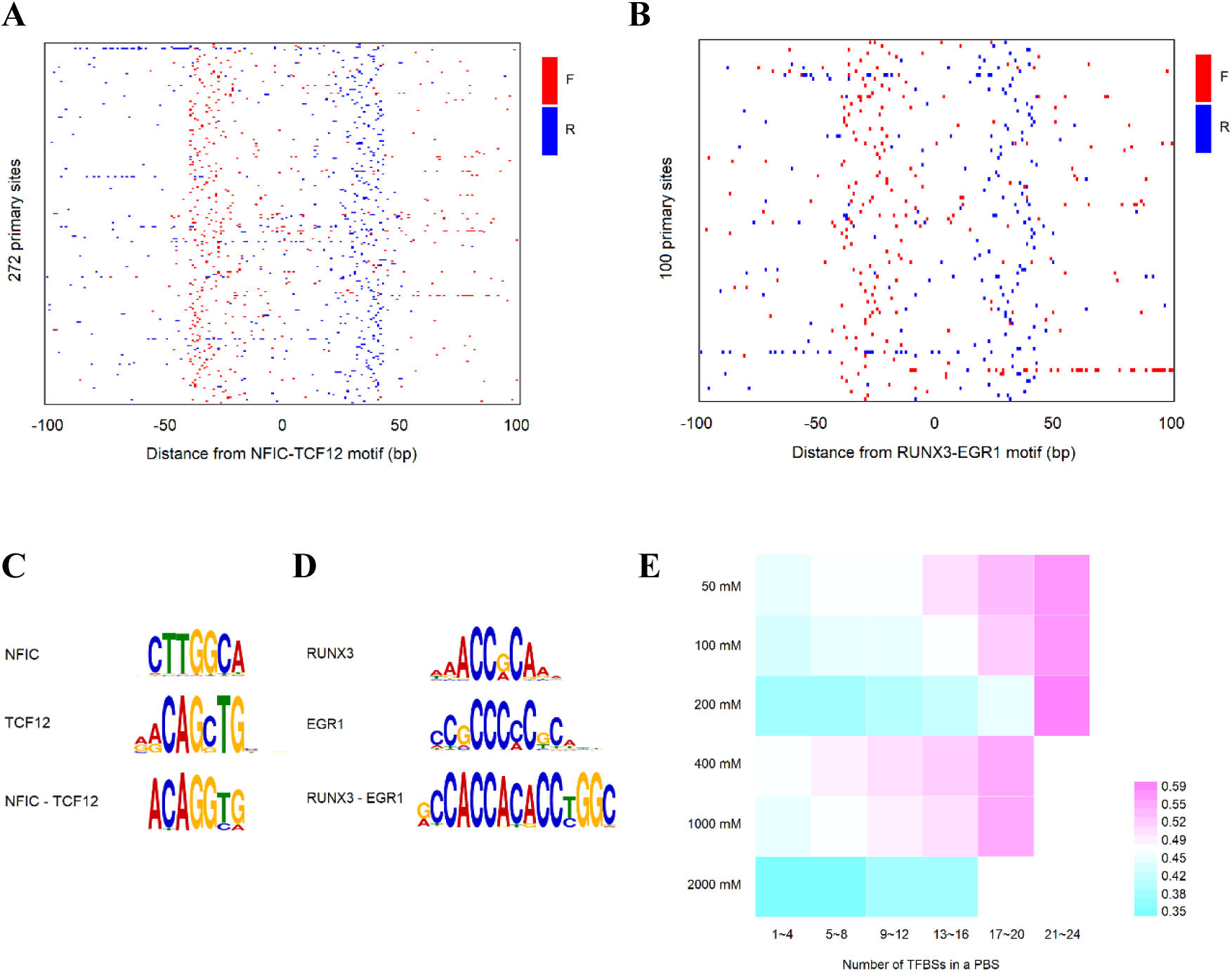
Co-localization of TFs predicated using PBSs. **(A-B)** Raw sequencing tag distribution around NFIC-TCF12-bound and RUNX3-EGR1-bound locations (rows). Red and blue represent the 5’ ends of forward (left border) and reverse strand tags (right border), respectively. **(C)** The sequences of NFIC-TCF12 recognition sites much like the two partially overlapped TF recognition sequences. **(D)** The sequences of RUNX3-EGR1 recognition sites much like the two partially overlapped TF recognition sequences. **(E)** The GC content affects the number of TFBSs contained in a PBS. PBSs with higher GC content tend to have more TFBSs. Purple indicates high GC content, and navy indicates low GC content.

### GC contents and lengths of TFBS-containing PBSs

We also investigated the GC content of PBSs. The results show that PBSs with high GC contents are more likely to bind with multiple TFs. A positive relationship exists between GC content and the number of TFBSs contained in the PBSs (Figure 4E). We further analyzed all predicted TFBSs and found that PBS sequences with high GC content (>50%) were more likely to bind multiple TFs or multiple TF families (Figure S6A). We then analyzed the correlation between number of TFBSs in PBSs and PBS length. The histograms (Figure S6B) showed a positive correlation between TFBS numbers and PBS lengths. The proportion of single TF-PBSs or PBSs containing single TF families decreased with increasing PBS length, indicating that longer PBSs are more likely to bind multiple TFs or TF-families. This suggests that sequence features and lengths of PBSs may contribute to TF binding strength. Longer PBSs with higher GC contents may bind multiple TFs and have stronger binding strengths than shorter PBSs with lower GC contents.

### Enrichment of PBSs containing TFBSs in the gene promoters

Next, we investigated the enrichment of the TFBSs in PBSs of promoter regions. We studied PBSs in the promoter regions and found that the content of TFBSs in the PBSs in the promoter region was higher than the average number of TFBSs in the PBSs genome-wide (Figure 5A-B). This confirms the notion that promoters tend to bind multiple TFs or TF families.

**Figure 5.**
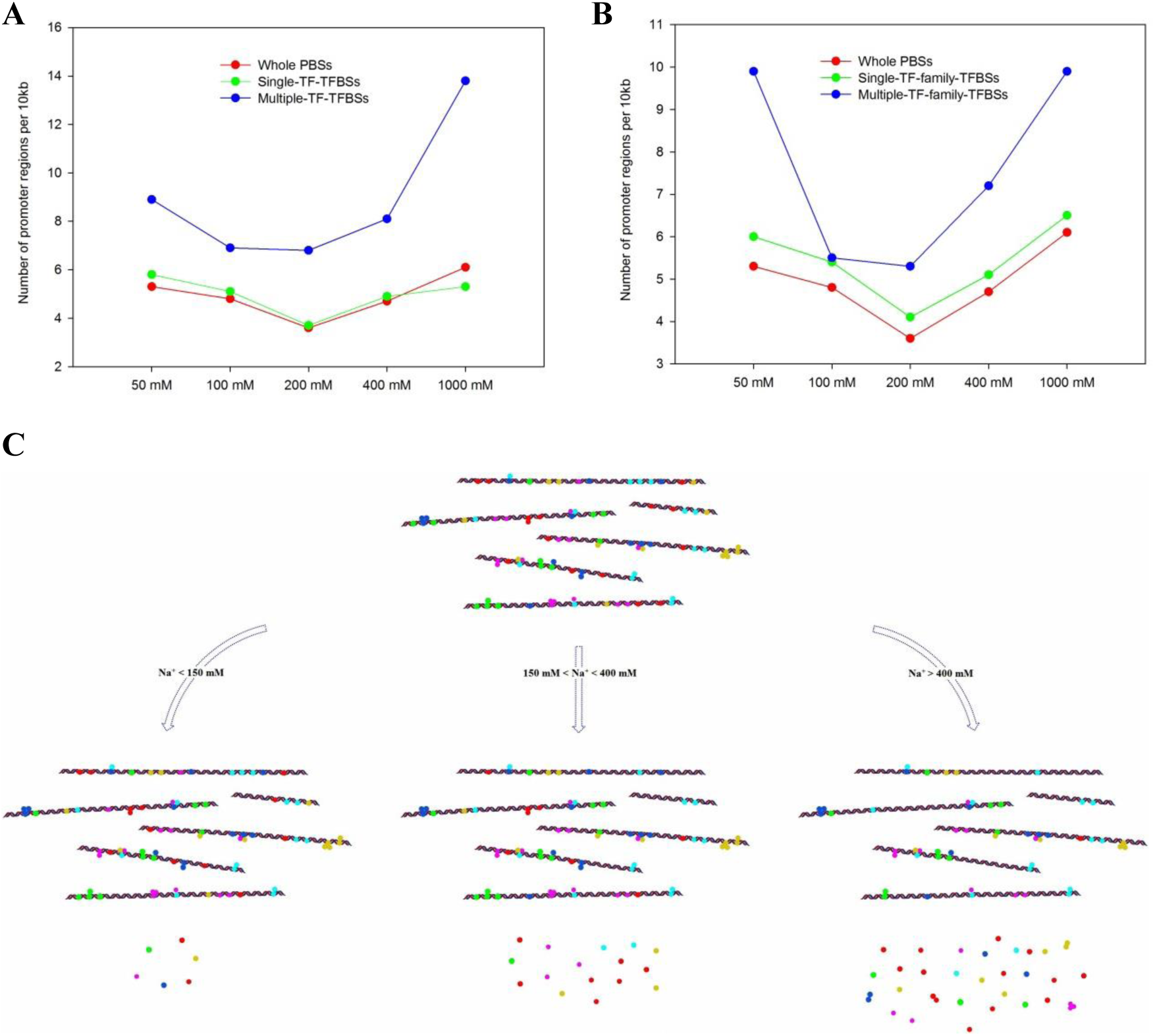
The number and type of TF binding to gene promoters varied with Na^+^ concentration. **(A)** PBSs containing multiple TFBSs are enriched in promoter regions. **(B)** PBSs containing binding sites for multiple TF families are highly enriched in promoters. **(C)** The model of TFs binding promoters in conditions of different Na^+^ wash environments. In the first case, a small number of TFs separate from promoters as Na^+^ concentration less than 150 mM. In the second, a larger proportion of active TFs break the combination with promoters as Na^+^ concentration in the range of 150 mM-400 mM. In the last, only single-TFs or TF complexes with strong strength to bind DNA, especially TF complexes consist of TFs derived from different TF-families, can still be remained on promoters as Na^+^ concentration greater than 400 mM.

We also observed different binding strengths of TFs to promoters. As shown in Figure 5A and 5B, the 200 mM NaCl concentration was a dividing point, suggesting that TFs associated with the 50 mM or 100 mM salt groups may have different regulatory functions than those associated with the 400 mM or 1000 mM salt groups.

Promoter PBSs were selected from 100mM and 1000mM salt PBS groups, and the genes corresponding to promoter were analyzed by GO enrichments. The genes in the 100 mM salt group were found to be enriched in molecular function such as protein serine/threonine kinase activity, cytoskeletal protein binding and GTPase activator activity (Figure S7A). The corresponding genes in the 1000 mM salt group were significantly enriched in molecular functions such as cell adhesion molecule activity, receptor activity and ubiquitin-specific protease activity (Figure S7B). A KEGG pathway enrichment analysis of the genes in the 100 mM group showed that they were enriched in signaling related to many types of growth factors (Figure S7C), which means that the TFs in this group mostly regulate positive cell growth process. The genes enriched with the 1000 mM salt group PBSs were enriched in biological pathways such as hypoxic and oxygen homeostasis (Figure S7D), which maintain basic cellular activities and inhibit over-exuberant life processes.

### Binding pattern and intensity of TFs to gene promoters

Since TFs primarily regulate gene expression by binding to gene promoters, the binding intensity is easy to be tested in the cases of Na^+^ washing environments. Based on our previous conclusions, we inferred the binding pattern of TFs to the gene promoters (Figure 5C). It is well known that the natural physiological salt concentration for cells is 150 mM NaCl. When the Na^+^ washing concentration is less than 150 mM, the binding between many TFs and DNA is less affected, so most TFs still bind to the promoters and continue to function. When the Na^+^ washing concentration is between 200 mM and 400 mM, some active TFs are separated from promoters due to their low tolerance to Na^+^. However, when the Na^+^ washing concentration increase above 400 mM, TFs that still binding to promoters show strong tolerance to high concentration of Na^+^, and this part of TFs tends to bind DNA in the form of TF complex to resist NaCl elution.

We also noticed that most of the TFBSs corresponding to promoters contained multiple motifs from different TF-families, which indicated that the binding strength between DNA and TFs depended not only on the number of TFs, but also on the type of TFs. It is speculated that TFs in the same family often have similar DNA binding domains (DBDs), resulting in the same or similar binding sites and a relatively single focus point on DNA. Therefore, TF complexes of the same family have relatively weak resistance to Na^+^. On the contrary, TFs of different families have different DBDs. When they bind DNA, TF complexes of different families will enhance their resistance to Na^+^ because they have more focus on DNA, so they can still bind DNA closely in high concentration Na^+^ washing environment. Figure 5C shows that the difference of eluted components under different concentrations of Na^+^ washing environment.

## Discussion

Protein-DNA interactions are essential components of all biological systems, fundamental to almost all biological processes, such as regulation of gene expression, DNA replication, repair, transcription, recombination, and packaging of chromosomal DNA. There are large number of DNA-binding proteins in gene regulatory regions, such as promoters and enhancers. However, the existing techniques, such as ChIP-seq, are not enough to capture most of protein binding sites in these specific regions at once. In this study, we describe a method called GWPBS-Cap for capturing genome-wide protein binding sites.

In the experiments, we optimized the method for crosslinking DNA-binding proteins and magnetic beads. We have tried to use EDC (1 - (3-dimethylaminopropyl) -3-ethylcarbodiimide hydrochloride) to mediate amino-carboxyl cross-linking between proteins and beads, but sequencing results showed that many non-specific short DNA fragments were enriched. This indicates that some Non-protein-binding DNA fragments cross-link with magnetic beads, probably use their exocyclic amino groups in the A, C, G nucleic acid bases. We then used sulfo-SMCC as cross-linking agent. The thiol groups of DNA binding proteins were cross-linked with amino groups in the magnetic beads. The use of sulfo-SMCC can effectively reduce the occurrence of false positive cross-links. In addition, there are many reductants, such as TCEP, DTT or beta-mercaptoethanol, which can be used to reduce disulfide bonds. The reason for choosing TCEP is not only that it has strong reduction capacity, but also that it does not contain sulfhydryl group and there is no risk of interfering with the follow-up reactions. Finally, after chromatin complexes were cross-linked with magnetic beads, cysteine is used to block amino sites on unbounded magnetic beads to minimize nonspecific DNA products.

A protein with stronger binding strength to DNA is usually resistant to high salt washing. In order to capture proteins with different binding strength to DNA, we washed the chromatin with buffers containing six different concentrations (50, 100, 200, 400, 1000, 2000 mM) of NaCl, respectively before crosslinking. PBSs were grouped according to the washing buffer used. We mapped those six groups of PBSs to the genome, including nucleosome sites and gene elements. The proteins in different eluents were also analyzed by protein Mass spectroscopy. Mass spectrometry confirmed that the content of TFs and other active proteins in 100-200 mM NaCl elution fractions was higher than that of other fractions. We also found that some histone variants were highly present in 50-100 mM NaCl elution fractions. It has been reported that some histone variants are unstable even in relatively low ionic conditions [15]. They are enriched near the transcription initiation sites (TSSs), which makes them sensitive to salt concentration [16].

Compared with other NaCl groups, the proportion of PBSs associated with promoter, 5’-UTR and 3’-UTRs was lower in 200 mM NaCl group, possibly because large number of proteins bound to these active regulatory regions were washed out at this NaCl concentration. In different salt groups, the proportion of PBSs corresponding to ORF and intergenic region is very high and stable, which indicates that in these regions, the content of various types of DNA-bound proteins is relatively high, and the distribution of different types of proteins was relatively balanced. This also indicates that the dynamics of DNA-protein interactions in these regions are different from those in promoters, 5’-UTRs and 3’-UTRs. Details of the interaction between DNA and protein in different regions need to be further explored.

TFs are DNA-binding proteins that play a key role in gene transcription. They are modular in structure and heterodimeric. Transcription factors bind to short conserved sequences located within the genome. We compared the PBSs sequence with the TF binding site data from ENCODE Consortium, and searched for the TFBSs in six groups of PBSs. The results showed that different TFs from the same family, or even from different TF families, could bind to the same PBSs, suggesting that multiple TFs interact in some local regions of the genome and play a regulatory role in the form of protein complexes. We found that the longer the PBS, the higher the GC content, and long PBSs tend to bind with more than one TFs.

For PBSs with multiple TFBS, this study can provide clues for how multiple TFS to cooperate with each other in a local genome region. Many TF binding sites across the genome have been identified by ChIP-seq, but do transcription factors that bind to the same region bind at the same time? The GWPBS-Cap method can help to understand whether multiple TFs bound to the same genome region also bind to the genome at the same time. If two TFs bound to the same genome region at the same time, it indicates that they may have collaborative effects. Moreover, through these clues, combined with information from databases such as STRING [17], the interactions between these transcription factor proteins could be identified, and how these TF interactions regulate local genome state can be studied. It has been reported that there are many TF combinations and collaborations in super enhancers [18] and other regions. GWPBS-Cap is a good complementary tool to ChIP-seq for genome research.

In addition, the function, localization and expression of TFBSs in promoter were analyzed. After comprehensive analysis, we found that PBSs in the promoter region of highly expressed genes tend to be less tolerant to high salt contents. On the contrary, PBSs in the promoter region of low-expression genes are more tolerant to high content salt washing and high ionic strength.

Our approach provides a tool for studying the protein binding strength in different regions of the genome and can be used to get overall feature of genome-wide DNA-protein interactions. In the future, single-cell GWPBS-Cap studies could shed light on the state of single-cell DNA-protein interactions. Genome-wide regulatory changes can be pursued by comparing DNA-protein interactions in single cells under different differentiation states or environmental stimuli.

In summary, GWPBS-Cap can be used to study the genome-wide distribution of protein binding sites without the use of antibodies. It could be used to obtain genome-wide information of the binding strength of various DNA-binding proteins in different genomic regions. Combine with TFBS databases, the method has potential to predict the TF-TF interactions in local genome regions. It will be a useful tool for studying chromatin structural dynamics and gene expression in differentiation and cancer.

## Materials and methods

### Cell culture

HeLa cells (from ATCC) were cultured in DMEM (HyClone, SH30243.01) supplemented with 10% FBS (Hyclone, SH30084.03) and 100 U ml^-1^ penicillin-streptomycin (HyClone, SV30010) and maintained in a incubator at 37°C and 5% CO_2_.

### GWPBS-Cap

#### Chromatin preparation

For each of the two GWPBS-Cap replicates in the different NaCl groups, 10^7^ HeLa cells were lysed by incubating with the cold Lysis Buffer (10 mM Hepes-KOH (pH 7.5), 1 mM EDTA, 10% glycerol, 0.5% NP-40, 0.25% Triton X-100 and protease inhibitor cocktail) at 4°C for 10 minutes. Nuclei were resuspended in 700 μl cold NaCl Buffer containing different concentrations (50 mM,100 mM, 200 mM, 400 mM, 1000 mM or 2000 mM) of NaCl, 10 mM Hepes-KOH (pH 7.5), 2 mM MgCl_2_, 2 mM EGTA and protease inhibitor cocktail. After rotation on a mechanical rotator at 4°C for 2 hours, nuclei were pelleted by centrifugation at 2000 rpm for 10 minutes. This step was repeated once and the pellet was resuspended in the same cold NaCl Buffer. Equal volumes of 2% formaldehyde solution (1% final concentration) were added to cross-link protein and DNA, followed by quench with 125 mM glycine. A Covaris S220 System (run at average amplitude for the shearing time course of 20 minutes in cold water) was used to shear the cross-linked DNA. Supernatants containing the solubilized chromatin DNA: protein complexes were clarified by spinning for 2 minutes at 13,200 rpm. TCEP (1:100 dilution of Thermo Scientific Bond-Breaker TCEP Solution, 77720) was added to a final concentration of 5 mM and the mixture incubated at room temperature for 30 minutes to reduce disulfide bonds of proteins, followed by two passes through a suitable desalting column. Ten percent of the chromatin sample was removed from each replicate for input sample preparation.

#### Covalent crosslinking

Following three washes with 0.01 M PBS (pH 7.4), 4.8 mg/ml Sulfo-SMCC (Thermo Fisher Scientific, 22322) was added and the samples were incubated on a mechanical rotator at room temperature for 30 minutes to activate M-270 Amine-coated Dynabeads (Thermo Fisher Scientific, 14307D). Excess cross-linkers were removed, and the activated beads were conjugated with the solubilized DNA: protein complexes on a mechanical rotator for 30 minutes at room temperature. Buffer containing 5 mM cysteine was added to stop the conjugation reaction.

#### Enzyme reactions and library preparation

Once covalently bound to the magnetic beads, the precipitated DNA samples were manipulated in the following order.

1. DNA end modification: 10 U T4 Polynucleotide Kinase (New England Biolabs, M0201) in 60 μl 1×T4 DNA Ligase reaction buffer were added to each sample and incubated at 37°C for 30 minutes.
2. Lambda exonuclease: 10 U λ exonuclease (New England Biolabs, M0262) in 60 μl 1×λ exonuclease reaction buffer was added and incubated at 37°C for 30 minutes.
3. Ligation of adapters to the 3’ ends of DNA fragments: Sp6-AS single-stranded adapter (5’ Phos-CTATAGTGTCACCTAAATCGTATG-NH_2_ 3’, synthesized by Sangon Biotech), was ligated to the DNA fragments by incubation in 60 μl 1×T4 RNA Ligase reaction buffer containing 1 mM ATP, 20% PEG8000 and 20 U T4 RNA Ligase 1 (New England Biolabs, M0204) at 16°C for 12 hours. In all cases, the beads were washed between each enzymatic step with the following buffers: a) Low Salt Wash Buffer (1% Triton X-100, 2 mM EDTA, 150 mM NaCl, 0.1% SDS, 20 mM Tris-HCl, pH 8.0), b) High Salt Wash Buffer (1% Triton X-100, 2 mM EDTA, 500 mM NaCl, 0.1% SDS, 20 mM Tris-HCl, pH 8.0), c) LiCl Wash Buffer (0.25 M LiCl, 1% NP-40, 1 mM EDTA, 1% Sodium deoxycholate, 10 mM Tris-HCl, pH 8.0), d) TE Buffer (1 mM EDTA, 10 mM Tris-HCl, pH 8.0).
4. Reversal of crosslinks and protein degradation: The chromatin samples were incubated with 200 μl proteinase K digestion buffer containing 30 μg Proteinase K (Roche), 0.2 M NaCl and 30 mM Tris-HCl at 65°C for 12 hours.
5. DNA extraction: The DNA was extracted with Phenol: Chloroform: Isoamyl alcohol (Amersco, 0883), precipitated with cold ethanol and dissolved in 11 μl water.
6. Preparation of double-stranded DNA: DNA samples were incubated with 1 μM Sp6 Primer (CATACGATTTAGGTGACACTATAG, oligos synthesized by Sangon Biotech), 3 μg BSA, 0.2 μM dNTPs in 20 μl 1×NEBuffer2 at 95°C for 5 minutes, then 55°C for 5 minutes. The samples were then cooled to room temperature, 5 U of Klenow Fragment (New England Biolabs, M0210) were added and incubated at 25°C for 20 minutes, followed by heat inactivation at 75°C for 20 minutes.
7. A-tail addition: DNA samples were incubated with 50 μM dATP and 5 U of Klenow Fragment (3’→5’ exo-) (New England Biolabs, M0212) in 30 μl 1 × NEBuffer2 at 37°C for 30 minutes.
8. Ligation of T7 adapter to the ends of double-stranded DNA: 0.5 μM of T7 a d a p t e r (5 ‘ O H -TA ATA C G A C T C A C TATA G G G A G A U - O H 3 ‘, 5 ‘ OH-TCTCCCTATAGTGAGTCGTATTA-NH_2_ 3’, synthesized and modified by Sangon Biotech) and 400 U T4 DNA ligase (New England Biolabs, M0202S) were added to a final volume of 40 μl in 1×T4 DNA Ligase reaction buffer, and incubated at 25°C for 60 minutes, followed by heat inactivation at 65°C for 10 minutes.
9. The DNA samples were purified with a NucleoSpin Gel and PCR Clean-up Kit (MACHEREY-NAGEL) following the manufacturer’s instructions.
10. PCR amplification and purification: PCR primers were synthesized by Sangon Biotech: T7-17 (GACTCACTATAGGGAGA), Sp6-19 (GATTTAGGTGACACTATA G). The PCR was done using a 20 μl PCR mix containing 2.5 U Pfu DNA Polymerase (a gift from Professor Shentao Li), 800 μM dNTPs, 1×GC buffer ? (Takara, 9155) for 30 PCR cycles. The PCR products were then purified with the NucleoSpin Gel and PCR Clean-up Kit.
11. Size selection and deep sequencing: PCR products with a length range between 80-120 bp were size selected on a 2% agarose gel, purified and DNA libraries were prepared. Paired-end sequencing (100 bp reads) was performed on an Illumina HiSeq 4000 platform by BGI Genomics (Wuhan, China).

### Protein identification with mass spectra

10^7^ HeLa cells were incubated in cold Lysis Buffer at 4°C for 10 minutes. Nuclei were then resuspended in 700 μl cold NaCl Buffer with different NaCl concentrations (50 mM, 50 mM, 100 mM, 200 mM, 400 mM, respectively). After rotation at 4°C for 2 hours, the nuclei were pelleted by centrifugation for 10 minutes at 2000 rpm. The supernatant was extracted with 50 mM NaCl buffer and saved. The nuclear proteins of the other four groups were extracted with 100 μl cold NaCl Buffer containing 100 mM, 200 mM, 400 mM, or 1000 mM NaCl, respectively. The samples were centrifuged for 10 minutes at 13,200 rpm, and supernatants were collected. After purification and trypsin digestion, the proteins were subjected to mass spectrometry analysis with a TripleTOF 6600 system (Sciex).

### RNA-seq

Total RNA was extracted from 3×10^6^ HeLa cells using TRIzol (Sigma, T9424). Prior to submission for high-throughput sequencing, an aliquot of the RNA from each sample was reverse transcribed using a RevertAid First Strand cDNA Synthesis Kit (Thermo Fisher Scientific, K1622). DNA library construction and sequencing were all performed by BGI Genomics. The libraries were sequenced with a 50bp single-end run on a BGISEQ 500 platform.

### SP1-ChIP-PCR

HeLa cells (10^7^ cells) were used to prepare a SP1-ChIP library. Sheared chromatin samples were incubated overnight with anti-SP1 antibodies (Santa Cruz Biotechnology, sc-17824X) conjugated to protein G-coated magnetic beads (ThermoFisher, 10003D). The cross-linking was reversed and the DNA was purified. PCR was performed and the products were purified with a NucleoSpin Gel and PCR Clean-up kit (MACHEREY-NAGEL).

The sequences of the chosen SP1-TFBSs and PCR primers are shown below.

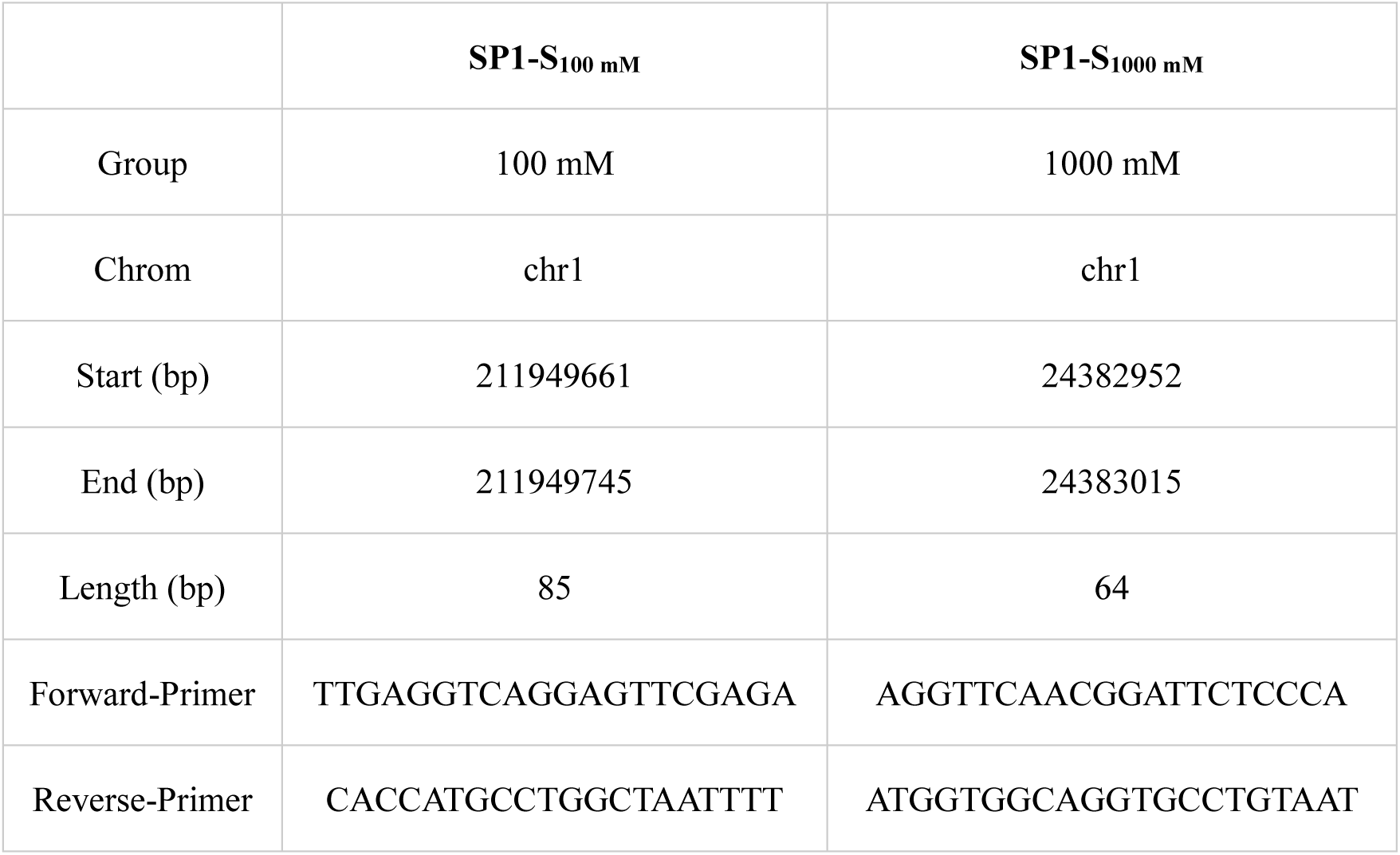

### Data analysis

#### Alignment to genomes and peak calling

The hg19 human genome reference sequence was obtained from UCSC Genome Browser (https://genome.ucsc.edu/cgi-bin/hgTracks?db=hg19&position=lastDbPos). The entire sequences of the GWPBS-Cap-seq reads, after removal of T7 adapters, which represent the fragments retained after λ exonuclease cleavage, were mapped to the reference genome using the SOAP2 aligner, allowing up to 2 mismatches. The resulting sequence read distributions were used to identify peaks. The peaks on forward (W) and reverse (C) strands were separated, the overlapping regions between the W and C strand peaks are considered to be PBSs.

#### TFBS library construction and prediction of TFBSs

The peak information for human TFBSs was downloaded from UCSC repository (http://hgdownload.cse.ucsc.edu/goldenPath/hg19/encodeDCC/, wgEncodeHaibTfbs, wgEncodeSydhTfbs, wgEncodeUwTfbs, wgEncodeRegTfbsClustered, wgEncode AwgTfbsUniform, wgEncodeUchicagoTfbs). The peak files of these six data sets were matched with our PBS sequences, and if a sequence in a PBS overlapped with a peak in the UCSC TF datasets, the sequence in a PBS was then defined as a TFBS.

#### Gene ontology

We used Gene Ontology (GO) enrichment analysis to map all target genes to GO terms with FUNRICH software [19], which can be used to calculate the gene or protein numbers for every GO term, and to plot the statistical results.

#### Motif discovery

MEME Suite [13] was used to find motifs on the PBSs.

#### DNase-seq, CTCF and histone modifications data

DNase-seq, CTCF and histone modifications data from HelaS3 cells were downloaded from the ENCODE Consortium datasets.

## Abbreviations

GWPBS-Cap: genome-wide protein binding site capture
PBS: protein binding site
TF: transcription factor
TFBS: transcription factor binding site
ChIP: chromatin immunoprecipitation
FPKM: fragments per kilobase of exon per million fragments mapped

## Data availability

Raw sequence reads and processed data from this study are available at NCBI Sequence Read Archive (SRA; http://www.ncbi.nlm.nih.gov/sra) under accession number SRP 152795 and the NCBI Gene Expression Omnibus (GEO; http://www.ncbi.nlm.nih.gov/geo) under accession number GSE116770, respectively.

## Authors’ contributions

YZ conceived the project. FL conducted the experiments and analyzed the data. TM contributed to the experiments. FL and YZ wrote the manuscript. All authors read and approved the final manuscript.

## Acknowledgements

We would like to thank Zhiyuan Xie and Shisheng Wang for bioinformatic support, and Haibo Zhang for helpful discussions. Sequencing was performed at BGI Genomics.

## References

1. Johnson DS, Mortazavi A, Myers RM, Wold B. Genome-wide mapping of in vivo protein-DNA interactions. Science. 2007; 316: 1497–1502.

2. Ren B, Robert F, Wyrick JJ, Aparicio O, Jennings EG, Simon I, et al. Genome-wide location and function of DNA binding proteins. Science. 2000; 290: 2306–9.

3. Sanders MM. Fractionation of nucleosomes by salt elution from micrococcal nuclease-digested nuclei. J. Cell Biol. 1978; 79: 97–109.

4. Teves SS, Henikoff S. Salt fractionation of nucleosomes for genome-wide profiling. Methods Mol Biol. 2012; 833: 421–32.

5. Henikoff S, Henikoff JG, Sakai A, Loeb GB, Ahmad K. Genome-wide profiling of salt fractions maps physical properties of chromatin. Genome Res. 2009; 19: 460–9.

6. Rhee HS, Pugh BF. Comprehensive genome-wide protein-DNA interactions detected at single-nucleotide resolution. Cell. 2011; 147: 1408–19.

7. Gerstein MB, Kundaje A, Hariharan M, Landt SG, Yan KK, Cheng C, et al. Architecture of the human regulatory network derived from ENCODE data. Nature. 2012; 489: 91–100.

8. Djebali S, Davis CA, Merkel A, Dobin A, Lassmann T, Mortazavi A, et al. Landscape of transcription in human cells. Nature. 2012; 489: 101–108.

9. Wingender E, Schoeps T, Haubrock M, Dönitz J. TFClass: a classification of human transcription factors and their rodent orthologs. Nucleic Acids Res. 2014; 43: 97–102.

10. Vaquerizas JM, Kummerfeld SK, Teichmann SA, Luscombe NM. A census of human transcription factors: function, expression and evolution. Nat Rev Genet. 2009; 10: 252–63.

11. Jolma A, Yin Y, Nitta KR, Dave K, Popov A, Taipale M, et al. DNA-dependent formation of transcription factor pairs alters their binding specificity. Nature. 2015; 527: 384–8.

12. Slattery M, Riley T, Liu P, Abe N, Gomez-Alcala P, Dror I, et al. Cofactor binding evokes latent differences in DNA binding specificity between Hox proteins. Cell. 2011; 147: 1270–82.

13. Bailey T, Boden M, Buske FA, Frith M, Grant CE, Clementi L, et al. “MEME SUITE: tools for motif discovery and searching”. Nucleic Acids Res. 2009; 37: 202–8.

14. Khan A, Fornes O, Stigliani A, Gheorghe M, Castro-Mondragon JA, van der Lee R, et al. JASPAR 2018: update of the open-access database of transcription factor binding profiles and its web framework. Nucleic Acids Res. 2018; 46: D1284.

15. Jin C, Felsenfeld G. Nucleosome stability mediated by histone variants H3.3 and H2A.Z. Genes Dev. 2007; 21: 1519–1529.

16. Jin C, Zang C, Wei G, Cui K, Peng W, Zhao K, et al. H3.3/H2A.Z double variant-containing nucleosomes mark ‘nucleosome-free regions’ of active promoters and other regulatory regions. Nat Genet. 2009; 41: 941–5.

17. Szklarczyk D, Morris JH, Cook H, Kuhn M, Wyder S, Simonovic M, et al. The STRING database in 2017: quality-controlled protein-protein association networks, made broadly accessible. Nucleic Acids Res. 2017; 45: 362–368.

18. Whyte WA, Orlando DA, Hnisz D, Abraham BJ, Lin CY, Kagey MH, et al. Master transcription factors and mediator establish super-enhancers at key cell identity genes. Cell. 2013; 153(2): 307–19.

19. Pathan M, Keerthikumar S, Chisanga D, Alessandro R, Ang CS, Askenase P, et al. A novel community driven software for functional enrichment analysis of extracellular vesicles data. J Extracell Vesicles. 2017; 6: 1321455.

